# Dr.seq2: a quality control and analysis pipeline for parallel single cell transcriptome and epigenome data

**DOI:** 10.1101/143271

**Authors:** Chengchen Zhao, Sheng’en Hu, Xiao Huo, Yong Zhang

## Abstract

An increasing number of single cell transcriptome and epigenome technologies, including single cell ATAC-seq (scATAC-seq), have been recently developed as powerful tools to analyze the features of many individual cells simultaneously. However, the methods and software were designed for one certain data type and only for single cell transcriptome data. A systematic approach for epigenome data and multiple types of transcriptome data is needed to control data quality and to perform cell-to-cell heterogeneity analysis on these ultra-high-dimensional transcriptome and epigenome datasets. Here we developed Dr.seq2, a Quality Control (QC) and analysis pipeline for multiple types of single cell transcriptome and epigenome data, including scATAC-seq and Drop-ChIP data. Application of this pipeline provides four groups of QC measurements and different analyses, including cell heterogeneity analysis. Dr.seq2 produced reliable results on published single cell transcriptome and epigenome datasets. Overall, Dr.seq2 is a systematic and comprehensive QC and analysis pipeline designed for parallel single cell transcriptome and epigenome data. Dr.seq2 is freely available at: http://www.tongji.edu.cn/~zhanglab/drseq2/ and https://github.com/ChengchenZhao/DrSeq2.

## Introduction

To better understand cell-to-cell variability, an increasing number of transcriptome technologies, such as Drop-seq [1, 2], Cyto-seq [3], 10x genomics [4], MARS-seq [5], and epigenome technologies, such as Drop-ChIP [6], single cell ATAC-seq (scATAC-seq) [7], have been developed in recent years. These technologies can easily provide a large amount of single cell transcriptome information or epigenome information at minimal cost, which makes it possible to perform analysis of cell heterogeneity on the transcriptome and epigenome levels, deconstruction of a cell population, and detection of rare cell populations. However, different single cell transcriptome technologies have their own features given their specific experimental design, such as cell sorting methods, RNA capture rates, and sequencing depths. But the methods and software such as Dr.seq [8] were developed for one single cell data type with certain functions (S1 File). Furthermore, the quality control step of single cell epigenome data is more challenging than for transcriptome data given the amplification noise caused by the limit number of DNA copy in single cell epigenome experiments. But few quality control and analysis method was developed specific for single cell epigenome data. Thus a comprehensive QC pipeline suitable for multiple types of single cell transcriptome data and epigenome data is urgently needed. Here, we provide Dr.seq2, a QC and analysis pipeline for multiple types of parallel single cell transcriptome and epigenome data, including recently published scATAC-seq data. Dr.seq2 can systematically generate specific QC, analyze, and visualize unsupervised cell clustering for multiple types of single cell data. For single cell transcriptome data, the QC steps of Dr.seq2 are primarily derived from Dr.seq [8] and the output of Dr.seq2 on these data will not be described in details in this paper.

## Materials and methods

### Drop-seq data

The Drop-seq samples were obtained from NCBI Gene Expression Omnibus (GEO) database under accession GSM1626793.

### MARS-seq data

The MARS-seq samples were obtained from NCBI Gene Expression Omnibus (GEO) database under accession GSE54006. These samples were combined as a MARS-seq dataset and analyzed by Dr.seq2 using three different dimension reduction methods.

### 10x genomics data

The 10x genomics datasets were obtained from 10x genomic data support (https://support.10xgenomics.com/single-cell/datasets). The sample named “50%: 50% Jurkat: 293T Cell Mixture” was analyzed by Dr.seq2 using three different dimension reduction methods.

### scATAC-seq data

The scATAC-seq datasets were obtained from NCBI Gene Expression Omnibus (GEO) database under accession GSE65360. We combined 288 scATAC datasets (GSM1596255 ~ GSM1596350, GSM1596735 ~ GSM1596830, GSM1597119 ~ GSM1597214) from three cell types and analyzed by Dr.seq2. Cell clustering was conducted for the combined scATAC-seq dataset. We also plotted the cell type labels using different colors on the clustering plot and found consistent classifications with the clustering results.

### Drop-ChIP data

The Drop-ChIP datasets were obtained from NCBI Gene Expression Omnibus (GEO) database under accession GSE70253.

### Implementation of Dr.seq2

Dr.seq2 was implemented using Python and R. Linux or MacOS environment with Python (version = 2.7) and R (version>=2.14.1) was suitable for Dr.seq2. It was distributed under the GNU General Public License version 3 (GPLv3). A detailed tutorial was provided on the Dr.seq2 webpage (http://www.tongji.edu.cn/~zhanglab/drseq2) and source code of Dr.seq2 was available on github (https://github.com/ChengchenZhao/DrSeq2).

### Quality control components

Dr.seq2 conducted four groups of QC measurements on single cell epigenome data: (i) reads level QC; (ii) bulk-cell level QC; (iii) individual-cell level QC; and (iv) cell-clustering level QC.

### Reads level QC and bulk-cell level QC

We used a published package called RseQC [9] for reads level QC of Drop-ChIP data and scATAC-seq data to measure the general sequence quality. In bulk-cell level QC, a Drop-ChIP dataset (or scATAC-seq datasets combined from several scATAC-seq samples) was regarded as a bulk-cell ChIP-seq (or bulk-cell ATAC-seq) data. Next, “combined peaks” were detected with total reads from the “bulk-cell” data using MACS[10] for output and the following steps. Different MACS parameters were applied to Drop-ChIP and scATAC-seq data. We used the published package CEAS to measure the performance of ChIP for ChIP-seq data (or Tn5 digestion for scATAC-seq data) [11].

### Individual-cell level QC

The reads number distribution was calculated by counting the number of reads assigned to each single cell. A single cell referred to a unique cell barcode in Drop-ChIP data. For scATAC-seq data, the peak number in each cell was defined as the number of “combined peaks” occupied by the reads in the cell. The distribution of different peak numbers in each cell indicated the different amount of information the cell contains.

### Cell-clustering level QC

Cells were first clustered based on their occupancy of “combined peaks” using hierarchical clustering. Next, cells in each cluster were regarded as the same cell type (or same cell sub-type), and reads from the same cell type were merged. For each cell type, unique peaks from other cell types were defined as specific peaks in this cell type. Specific peaks in different cell types were displayed with different colors according to genomic locations. Silhouette method is used to interpret and validate the consistency within clusters defined in previous steps.

Note that reads with no overlap with “combined peaks” were discarded in this step and the following steps. Clusters containing less than 3 single cells were also discarded.

### Simulation of scATAC-seq datasets

To measure the tolerance of Dr.seq2 for low sequencing depth and small numbers of cells of a certain cell type, we simulated datasets from 3 cell types with different cell proportions and sequencing depths using scATAC-seq data (Table 1). To test the effect of low sequencing depth, we sampled the reads count from 10,000 reads to 100,000 reads for each cell and compared these results with the Goodman-Kruskal’s lambda index [12] of clustering results using cells with a certain number of reads.

**Table 1.** Meta data and accession ID for the scATAC-seq data used in simulation for pipeline tolerance evaluation.

| Accession ID | Cell line | Cell type | Target/regular cell |
| --- | --- | --- | --- |
| GSM1596255~GSM1596350 | H1 | human embryonic stem cell line | Target |
| GSM1596735~GSM1596830 | GM12878 | lymphoblastoid cells | Regular |
| GSM1597119~GSM1597214 | K562 | chronic myeloid leukemia cells | Regular |

We defined the 1 out of 3 cell types as “target cell type”, while the other cell types were defined as “regular cell type”.

To test the effect of low cell numbers of a certain cell type (defined as a target cell type) on cell clustering, we defined 1 of the 3 cell types as the “target cell type”, whereas the other cell types were defined as the “regular cell type”, and sampled cells with following compositions: 10:70:70 (10 for target cell type, 70 for the two regular cell types), 15:67:67, 20:65:65, 25:62:62, 30:60:60, 35:57:57, 40:55:55, 45:52:52 and 50:50:50. Then, we called “combined peaks” and clustered cells on the simulated dataset. The Goodman-Kruskal’s lambda index [12] was calculated to evaluate the cell clustering performance. The average Goodman-Kruskal’s lambda index and 95% confidence intervals were calculated from 20 simulations.

## Results and discussion

### Dr.seq2 overview

The Dr.seq2 QC and analysis pipeline is suitable for both single cell transcriptome data and epigenome data. Multiple types of single cell transcriptome data (including scRNA-seq, Drop-seq, inDrop, MARS-seq and 10x genomics data) and epigenome data (including scATAC-seq and Drop-ChIP) are acceptable for Dr.seq2 with relevant functions (S1 Fig).

Recently many methods and software were developed for single cell RNA-seq data. However most of them were suitable for certain data types with limited functions. We compared the major function of Dr.seq2 to existing state-of-the-art methods (Table 2). Dr.seq2 provides two advantages: 1) Dr.seq2 supports different types of single cell transcriptome data and single cell epigenome data. 2) Dr.seq2 provides both multifaceted QC reports and cell clustering results. Then We used the simulated single cell RNA-seq data from seven RNA-seq datasets from ENCODE (S2 File) to estimate the performance of our Dr.seq2 pipeline (using different dimensional reduction methods: SIMLR and t-SNE) in cell clustering comparing to three existing methods (SINCERA, SNN-Cliq, BackSPIN). We applied these five methods on ten datasets with different numbers of reads per cell range from 100 to 10,000 to measure the accuracy and time cost of each method on different sequencing depth. SIMLR shows more accurate clustering results than t-SNE on the datasets with small number of reads per cell and comparable clustering results on the datasets with large number of reads per cell. And Dr.seq2 (using either SIMLR or t-SNE) shows better clustering accuracy than SNN-Cliq, and comparable clustering accuracy with BackSPIN and SINCERA on the datasets with large number of reads per cell. On the datasets with small number of reads per cell, SINCERA clustering result shows better accuracy than Dr.seq2 (using either SIMLR or t-SNE) and SNN-Cliq. However SINCERA takes a great mount of time on all these datasets comparing with Dr.seq2. As for BackSPIN, it does not support for these datasets with small number of reads per cell. Overall, Dr.seq2 (using either SIMLR or t-SNE) provides reliable cell clustering results with acceptable time cost (S2 Fig).

**Table 2.**
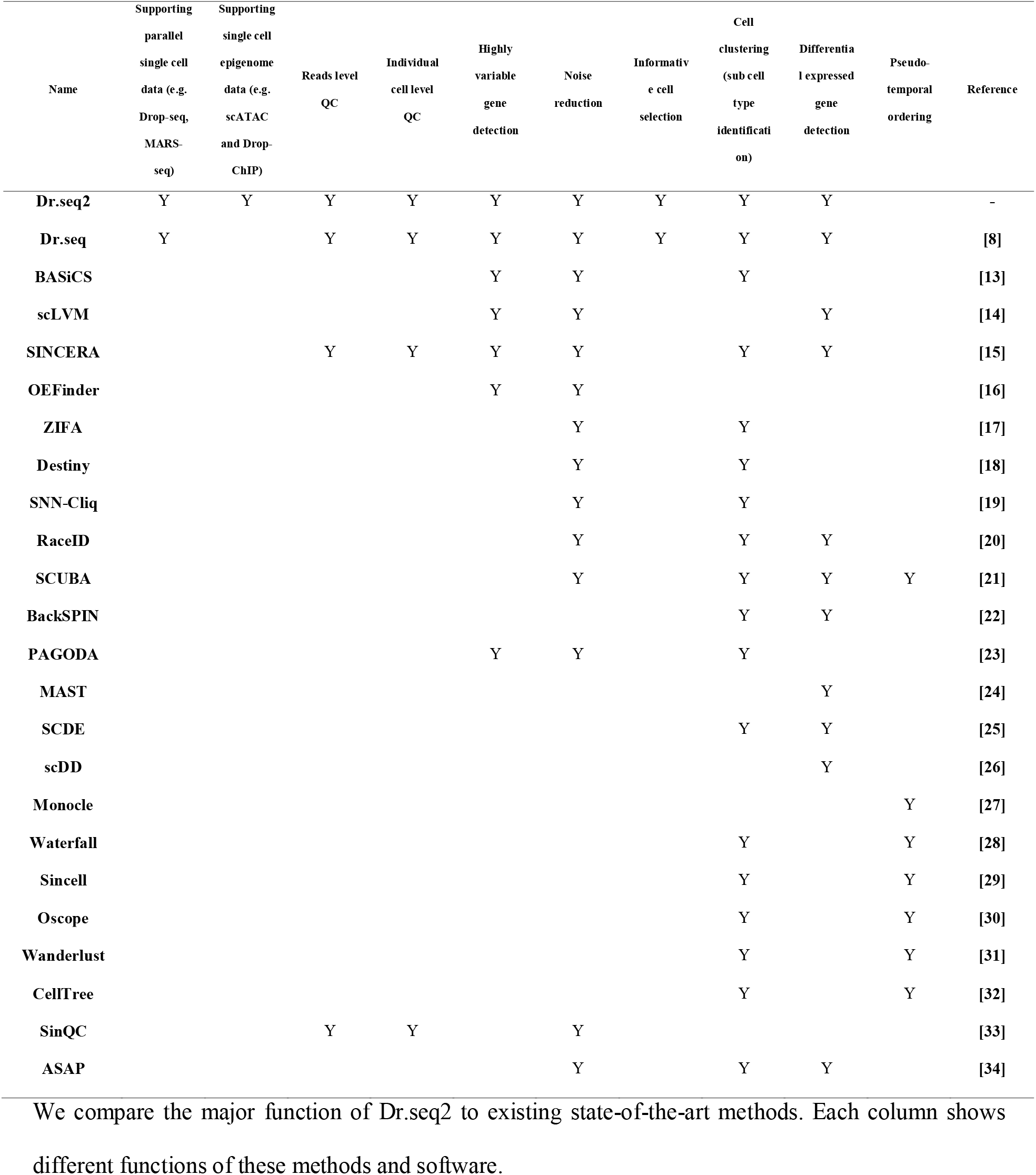
Comparison of functions between Dr.seq2 and other software developed for single cell transcriptome data.

### QC and analysis workflow

Dr.seq2 uses raw sequencing files in FASTQ format or alignment results in SAM/BAM format as input with relevant commands and generates four steps of QC measurements and analysis results (Fig 1).

**Fig 1.**
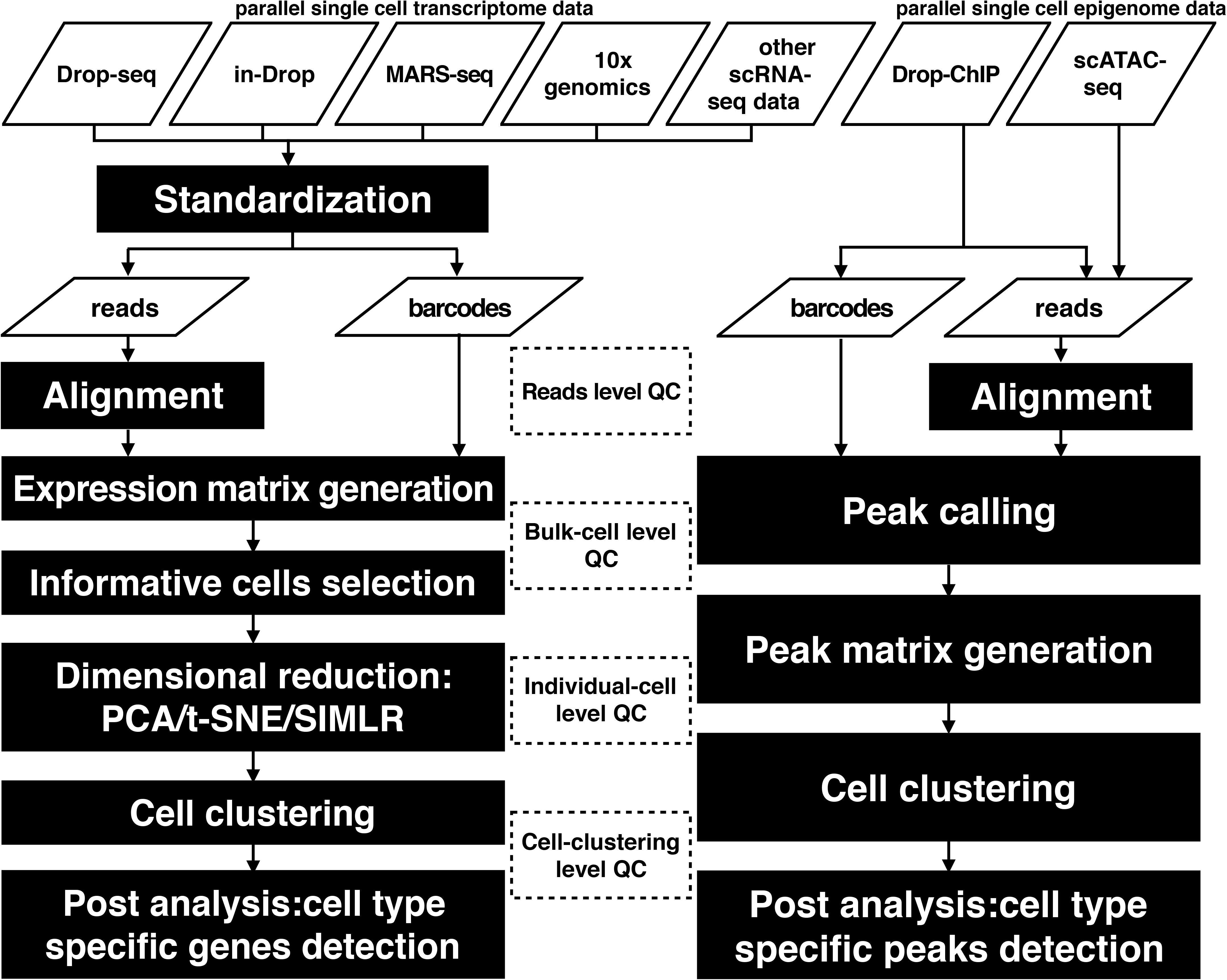
Flowchart illustrating the Dr.seq2 pipeline with default parameters. The workflow of the Dr.seq2 pipeline includes QC and analysis components for parallel single cell transcriptome and epigenome data. The QC component contains reads level, bulk-cell level, individual-cell level and cell-clustering level QC.

For transcriptome data, the QC steps of Dr.seq2 are primarily derived from Dr.seq [8]. However, almost all data types are now supported, and more dimension reduction methods, including PCA, t-SNE and SIMLR[35], are supported. For single cell epigenome data, technologies like scATAC-seq and Drop-ChIP are increasingly common. However few quality control and analysis approaches have been developed for these data. Dr.seq2 conducts QC measurements on single cell epigenome data from four aspects: (i) reads level QC, including sequence quality, nucleotide composition and GC content of reads inherited from previous work; (ii) bulk-cell level QC, including genomic distribution of “combined peaks” and average profile on regulatory regions; (iii) individual-cell level QC, including the distribution of the number of reads and the peak number distribution; and (iv) cell-clustering level QC, including Silhouette score[36] and cell type-specific peak detection.

### Cell clustering for different single cell transcriptome data types using different dimension reduction methods

We applied our pipeline to three different types of single cell transcriptome data (Drop-seq, MARS-seq and 10x genomics data) using three different dimension reduction methods (PCA, t-SNE and SIMLR[35]) to evaluate the performance of Dr.seq2 on different types of single cell transcriptome data (Fig 2). Due to the different distance calculation method and kernel function the method used, Dr.seq2 represented cluster results from different dimensions.

**Fig 2.**
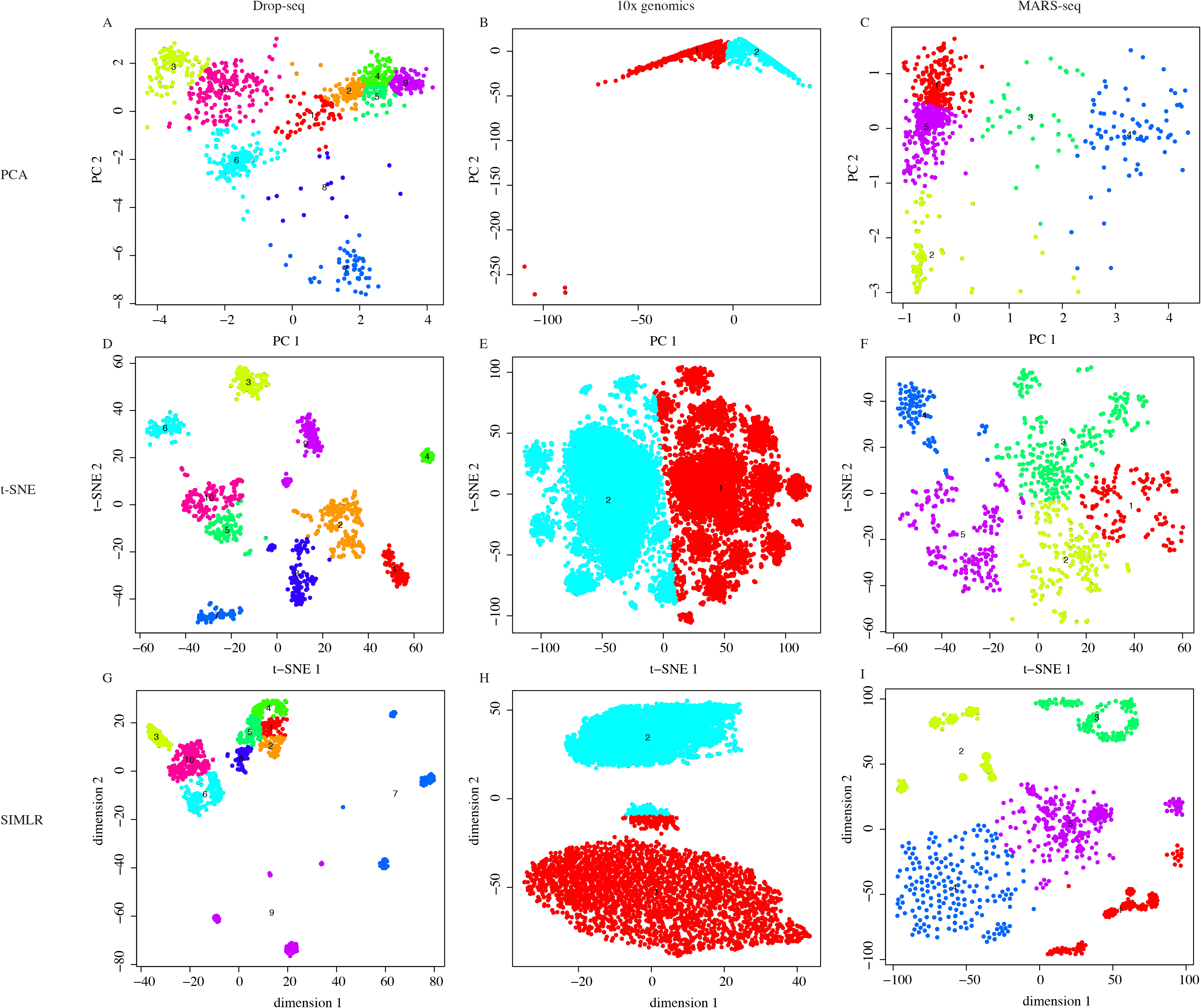
Dimensional reduction results for different single cell transcriptome data types. (A-I) Cell clustering results using dimensional reduction methods (PCA, t-SNE and SIMLR) on different types of single cell transcriptome data (Drop-seq, 10x genomics and MARS-seq).

### Bulk-cell level QC of scATAC-seq data to measure the performance of Tn5 digestion

To evaluate the performance of Dr.seq2 on single cell epigenome data, we combined 288 scATAC datasets (GSM1596255 ~ GSM1596350, GSM1596735 ~ GSM1596830, GSM1597119 ~ GSM1597214) from three cell types and applied Dr.seq2 to it. “Combined peaks” were detected with total reads from the combined dataset using MACS for output and the following steps. We measured the scATAC data quality in bulk-cell level from 4 aspects (Fig 3): 1) Peak distribution on each chromosome; 2) Open regions distributed over the genome along with their scores; 3) Average profiling on different genomic features; 4) Fragment length distribution. The peak distribution on each chromosome and the open region distributed over the genome showed the general quality of Tn5 digestion. The average profiling on different genomic features represented the quality of Tn5 digestion around specific regions.

**Fig 3.**
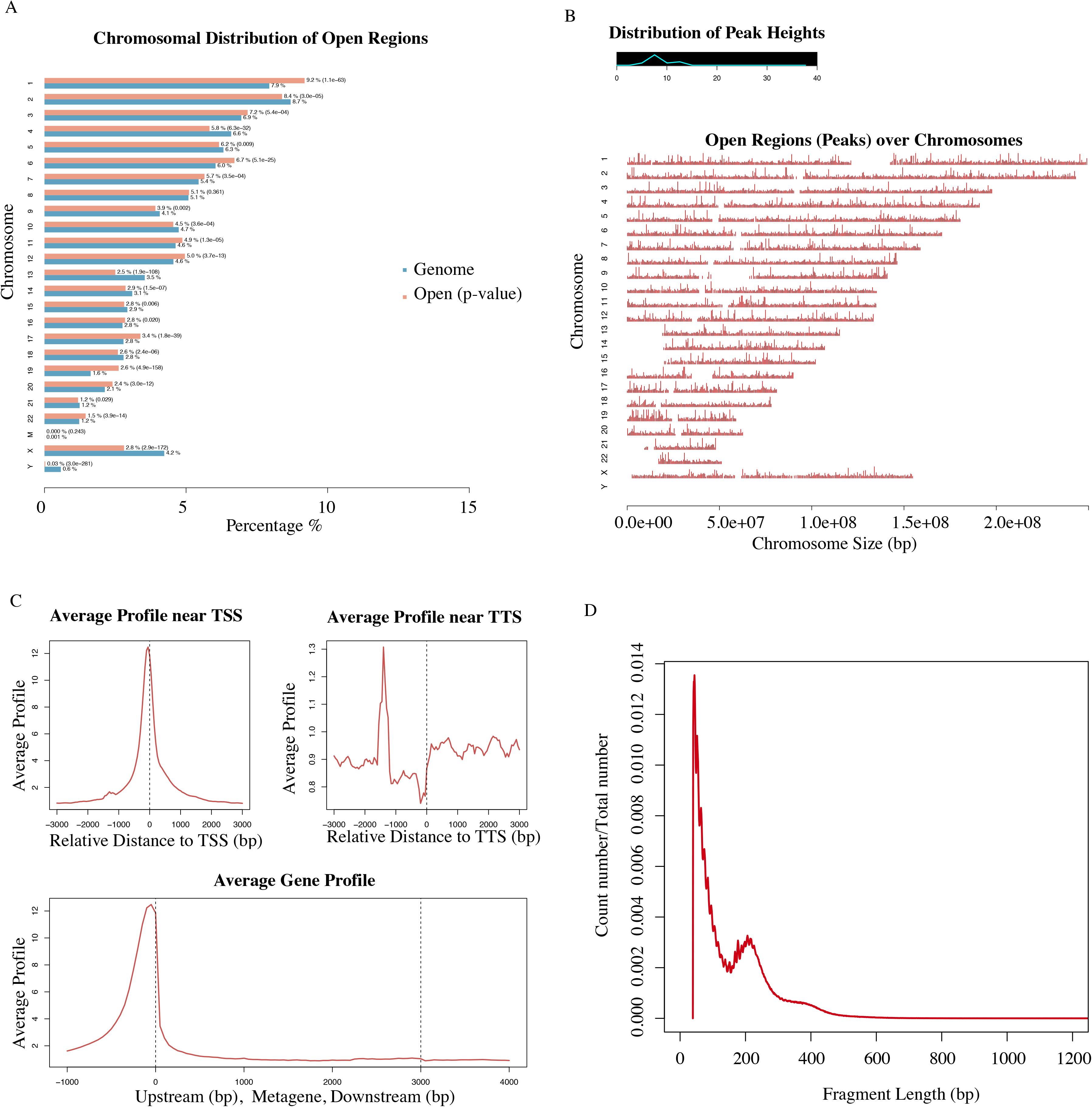
Bulk-cell level QC for scATAC-seq datasets. **A)** Peak region number distribution on each chromosome. The blue bars represent the percentages of the whole tiled or mappable regions in the chromosomes (genome background) and the red bars showed the percentages of the whole open region. These percentages are also marked right next to the bars. P-values for the significance of the relative enrichment of open regions with respect to the gnome background are shown in parentheses next to the percentages of the red bars. **B)** Open region distribution over the genome along with their scores or peak heights. The line graph on the top left corner illustrates the distribution of peak score. The x-axis of the main plot represents the actual chromosome sizes. **C)** Average profiling on different genomic features. The panels on the first row display the average enrichment signals around TSS and TTS of genes, respectively. The bottom panel represents the average signals on the meta-gene of 5 kb. **D)** Red line shows number distribution of different fragment length.

And the periodicity fragment length distribution indicated factor occupancy and nucleosome positions due to different Tn5 digestion degrees.

### Cell clustering for scATAC-seq datasets with three clusters that were consistent with the cell type labels

To measure the cell clustering performance of Dr.seq2 on epigenome data, cells from the combined scATAC-seq dataset were firstly clustered based on their occupancy of “combined peaks” using hierarchical clustering. Then cell type labels were marked by different colors according to the original cell type information (red stand for H1 cells, yellow stand for GM12878 cells and blue stand for K562 cells). Cells were clearly separated into different groups that were consistent with the cell type labels by Dr.seq2 (Fig 4A).

**Fig 4.**
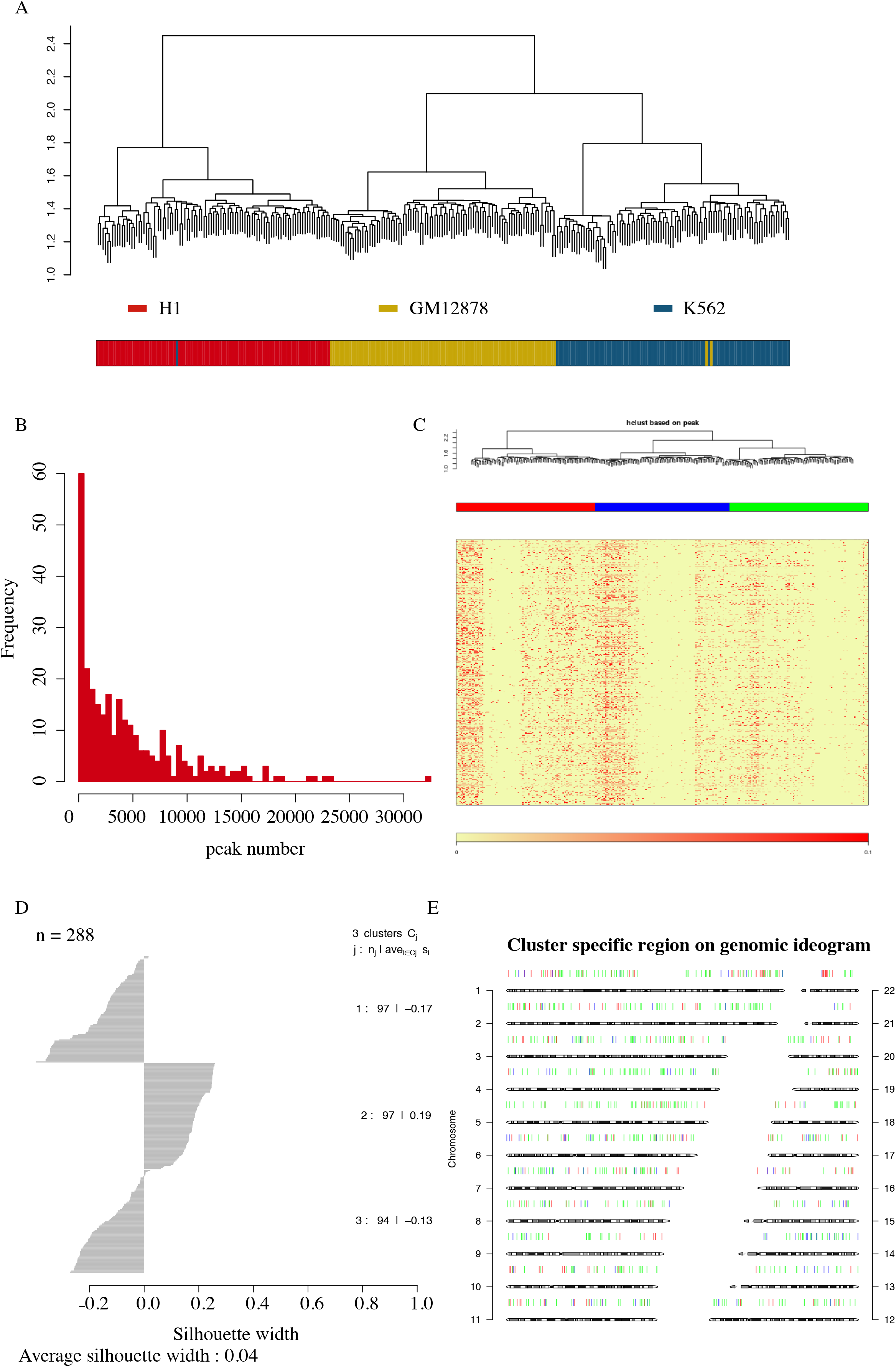
Cell-clustering level QC and single-cell level QC for scATAC-seq data. A) Upper panel shows cell-clustering results for combined scATAC samples generated from 3 different cell types. Bottom panel shows corresponding cell type labels of each cell marked by different colors (red stand for H1 cells, yellow stand for GM12878 cells and blue stand for K562 cells). The clustering step of Dr.seq2 clearly separated the scATAC-seq samples from three different cell types into different groups that were consistent with the cell type labels. B) Distribution of peak number for each single cell. C) Cell Clustering tree and peak region in each cell. The upper panel represents the hieratical clustering results based on each single cell. The second panel with different colors represents decision of cell clustering. The bottom two panels (heatmap and color bar) represent the “combined peaks” occupancy of each single cell. D) Barplot shows Silhouette score of each cluster. Silhouette method is used to interpret and validate the consistency within clusters defined in previous steps. E) Cluster specific regions in each chromosome. Specific regions for different cell clusters are marked by different colors and ordered according to genomic loci.

### Single-cell level QC and post analysis of scATAC-seq data

In the single-cell level QC of Dr.seq2 on scATAC-seq data, the peak number of in each cell was defined as the number of “combined peaks” occupied by the reads in the cell. The distribution of different peak numbers in each cell indicated the different amount of information the cell contains (Fig 4B). Cell clustering was conducted based on the peak information in each cell using hierarchical clustering and open region was shown in the order of genomic location (Fig 4C). And Silhouette score [36] validated the consistency of each cluster (Fig 4D). Then cells in the same clusters were considered as cells in the same cell type and combined for the detection of cell type specific regions, which were defined as the peak regions that only covered in this cell type. Specific regions for different cell clusters were marked by different colors and ordered according to genomic loci (Fig 4E).

### Cell clustering stability on simulated scATAC-seq data

To measure the tolerance of Dr.seq2 for low sequencing depth and small numbers of cells of a certain cell type, we simulated datasets with different cell proportions and sequencing depths by using scATAC-seq datasets from three cell types (Table 1). We selected cells in different proportion with 100,000 reads per cell and then performed cell clustering using Dr.seq2. The performance of cell clustering methods was evaluated by Goodman-Kruskal’s lambda index. And the average Goodman-Kruskal’s lambda index calculated from 20 simulations indicated that Dr.seq2 was suitable for cell clustering with different cell proportions (Fig 5A). We also selected fifty cells from each cell type with the reads count range from 10,000 reads to 100,000 reads for each cell to measure the tolerance of Dr.seq2 on low sequence depth. Dr.seq2 produced stable clustering results with greater than 40,000 reads per cell (Fig 5B).

**Fig 5.**
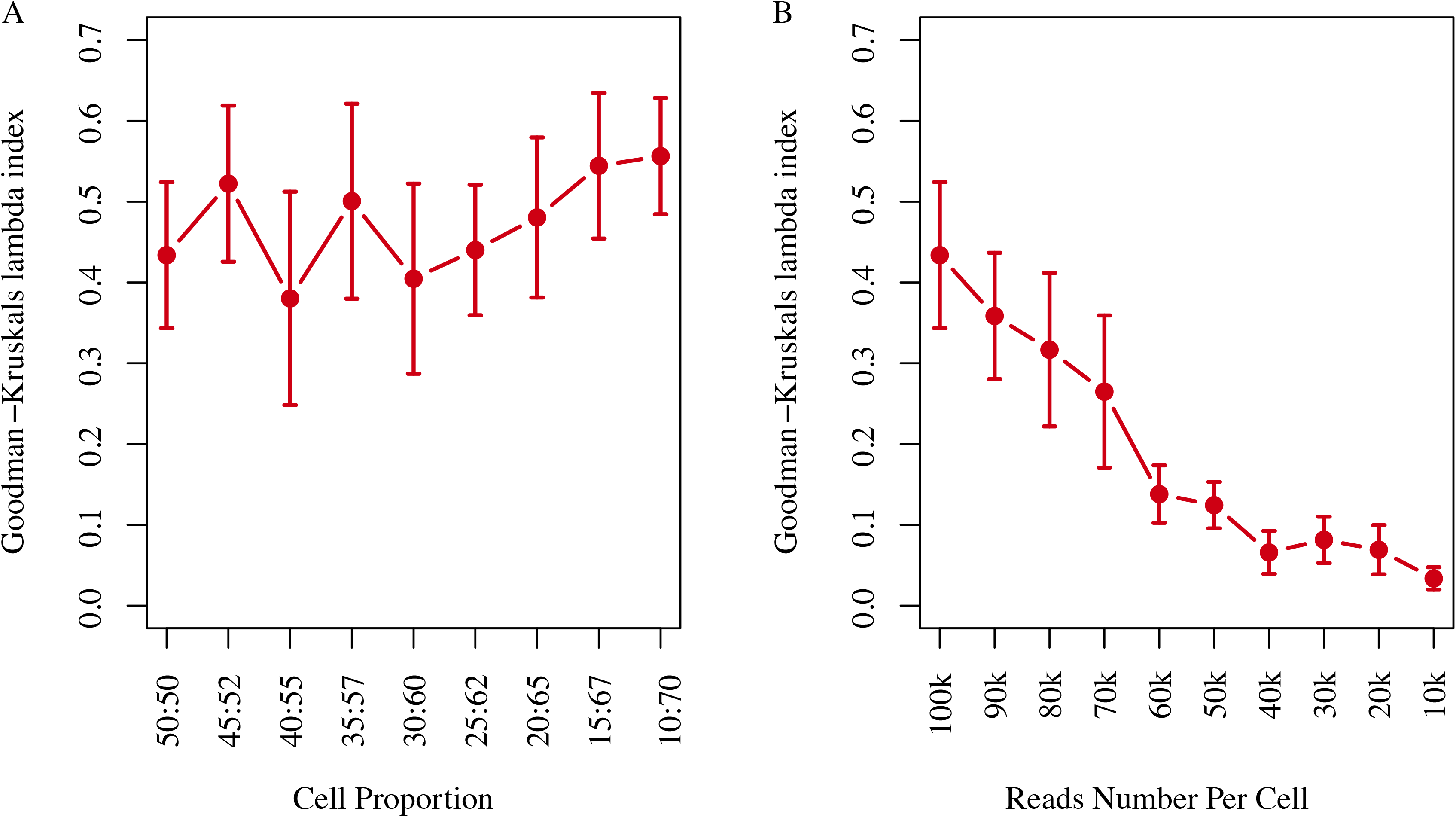
Cell clustering stability on simulated scATAC-seq data. **A)** Clustering stability of Dr.seq2 on simulated data with different numbers of reads per cell. The lambda index (y-axis) is plotted as a function of the number of reads per cell (x-axis). Error bars represent 95% confidence intervals calculated from 20 simulations. **B)** Clustering stability of Dr.seq2 on simulated data with different cell proportion depths. The lambda index (y-axis) is plotted as a function of the target cell number (x-axis). Error bars represent 95% confidence intervals calculated from 20 simulations.

### Computational cost of Dr.seq2

We also measured the computational time cost of Dr.seq2 by applied Dr.seq2 on combined scATAC-seq datasets (Table 3). The running time of each step was calculated using a single CPU (Intel® Xeon® CPU E5-2640 v2 @ 2.00 GHz).

**Table 3.** Running time of each QC and analysis step for scATAC datasets.

| Steps | Time (s/CPU) | Percentage (%) |
| --- | --- | --- |
| Merge Cells | 1507 | 39.72 |
| Bulk-cell level QC | 1654 | 43.60 |
| Individual-cell level QC and cell-clustering QC | 626 | 16.50 |
| Post-analysis | 5 | 0.13 |
| Summary Report | 2 | 0.05 |

288 scATAC datasets from three cell types were used to evaluate the runtime of Dr.seq2. The running time for each step was calculated using a single CPU (Intel® Xeon® CPU E5-2640 v2 @ 2.00 GHz).

## Conclusions

In summary, Dr.seq2 is designed for QC and analysis components of parallel single cell transcriptome and epigenome data. Parallel single cell transcriptome data generated by different technologies can be transformed to the standard input for Dr.seq2 with contained functions. Using relevant commands, Dr.seq2 can also be used to report quality measurements based on four aspects and generate detailed analysis results for scATAC-seq and Drop-ChIP datasets.

## Acknowledgments

We thank Shiyang Zeng and Yiying Lang for their suggestions.

## Supporting information

**S1 Fig. Workflow displays the software structure and detailed QC steps of Dr.seq2. A)** Dr.seq2 provides QC and analysis for three major data types: single cell transcriptome data (DrSeq part), Drop-ChIP data (DrChIP part) and scATAC-seq data (ATAC part). For single cell RNA-seq data, two additional step-by-step functions are included: 1. Expression matrix generation for amounts of single cell RNA-seq datasets (GeMa step) and 2. Cell clustering and analysis for the single cell expression matrix (comCluster step). For different parallel single cell RNA-seq technologies, input data are standardized for DrSeq part. **B)** Four groups of QC measurements are conducted on single cell transcriptome data and epigenome data: 1.Reads level QC including reads quality, reads nucleotide composition and reads GC content 2.Bulk-cell level QC including reads alignment summary and gene body coverage for transcriptome data; peak distribution; average profile on regulatory region and the distribution of different numbers of fragment length for epigenome data. 3. Individual-cell level QC including duplicate rate distribution, covered gene number and intron rate distribution and intron rate distribution for transcriptome data; peak number distribution and fragment length distribution for epigenome data. 4. Cell-clustering level QC including Gap statistics score and Silhouette score for transcriptome data, h-clustering and cluster specific peaks for epigenome data.

**S2 Fig. Comparing the performance of Dr.seq2 and three existing state-of-the art methods on cell clustering. A)** Clustering accuracy measured by the Goodman-Kruskal’s lambda index of Dr.seq2 t-SNE, Dr.seq2 SIMLR methods and three published methods on simulated data with different numbers of reads per cell. The lambda index (y-axis) is plotted as a function of the number of reads per cell (x-axis). **B)** Running time of Dr.seq2 t-SNE, Dr.seq2 SIMLR methods and three published methods on simulated data with different numbers of reads per cell. The running time (y-axis) is plotted as a function of the number of reads per cell (x-axis). The running time for each method was calculated using a single CPU (Intel® Xeon® CPU E5-2640 v2 @ 2.00 GHz).

**S1 File. Comparison of functions between Dr.seq2 and other software developed for single cell transcriptome data.**

**S2 File. Meta data and accession ID for the bulk-cell RNA-seq data used in simulation.**

**S3 File. Dr.seq2 QC and analysis output report for the scATAC-seq dataset.**

**S4 File. Dr.seq2 QC and analysis output report for the Drop-ChIP dataset.**

**S5 File. Dr.seq2 QC and analysis output report for the 10x genomics dataset.**

